# Sustained Delivery of a Shingles Subunit Vaccine Overcomes Age-Related Declines in Humoral and Cellular Immunity Relative to Shingrix

**DOI:** 10.64898/2026.03.11.711122

**Authors:** Ye Eun Song, Jerry Yan, Ben S. Ou, Olivia M. Saouaf, Noah Eckman, Eric A. Appel

## Abstract

The global aging population faces heightened vulnerability to infectious diseases due to immunosenescence, which diminishes the potency, durability, and breadth of vaccine-induced immunity. While the leading shingles vaccine, Shingrix®, provides protection against herpes zoster virus for adults aged 50 and older, it is often associated with severe local and systemic reactogenicity, which limits vaccine compliance. Here, we report an injectable polymer-nanoparticle (PNP) hydrogel platform for the sustained delivery of a shingles subunit vaccine to enhance immune responses while mitigating reactogenicity in aged mice. Hydrogel-based vaccination elicited significantly more potent and durable humoral immune responses than Shingrix®, while inflammatory cytokine levels remained below the limit of detection. Moreover, aged mice vaccinated with the hydrogel-based vaccine exhibited robust antigen-specific cellular immune responses. These findings demonstrate that controlling the temporal presentation of vaccine components can overcome age-associated declines in immune responsiveness without inducing excessive inflammatory signaling. By decoupling immunogenicity from reactogenicity, our hydrogel-based delivery strategy offers a promising approach to improve both the efficacy and tolerability of subunit vaccines for the elderly and may be broadly applicable to other vaccines targeting aging populations.

## 1. Introduction

The proportion of elderly people (over 60 years of age) in the global population is increasing, and this trend is expected to continue for many years.^[1]^ An aging society presents significant challenges to public health due to the frequent infections and poor outcomes from severe infection in the elderly.^[2, 3]^ While vaccination is a crucial tool for preventing infectious diseases, elderly populations also suffer from reduced vaccine efficacy due to age-related immunosenescence.^[4–6]^ Indeed, in the case of influenza, which is listed as top ten leading cause of death in elderly, vaccine efficacy is only around 30-50% in people over 65 years-old compared to 70-90% in young people.^[7]^ Efforts to improve vaccine design for the elderly are ongoing with a primary focus on increasing vaccine potency by increasing the dosage of the vaccine components and/or incorporating novel adjuvants to increase immune activation.^[8]^ For example, AS01 (GSK) is a liposome-based adjuvant containing two immunostimulants, including monosphosphoryl lipid A (MPLA) and saponin (QS-21), that is used in the Shingrix® (GSK) subunit protein vaccine for varicella herpes zoster virus (VZV). Preclinical rodent studies showed that AS01 elicited the highest gE-specific humoral and cellular immune responses compared to other adjuvants, including alum, oil-based emulsions (AS02; GSK), or a mixture of liposomes comprising either MPLA or QS21.^[9]^ Two phase 3 clinical trials demonstrated over 90% efficacy in immunocompetent adults over 50 years old, and Shingrix® is currently licensed for the prevention of shingles in many countries worldwide.^[10, 11]^ Despite success in protection against disease, Shingrix® suffers from low vaccine acceptance since AS01 is often associated with strong reactogenicity and adverse reactions both locally at the injection site and systemically (*e.g.*, fever, headache, and fatigue).^[10–15]^ Although adverse symptoms are not prolonged, concerns regarding vaccine tolerability have impacted patient compliance and resulted in lower vaccine coverage compared to vaccines for other indications recommended for aged adults (*e.g.*, influenza or pneumonia).^[15, 16]^ Despite the recent advancements in adjuvants, new technologies are still needed to elicit potent and durable immune responses in aged people with minimal reactogenicity and negligible undesirable side effects.

Recently, prolonged vaccine exposure has demonstrated the ability to improve humoral and cellular immune responses.^[17–20]^ Our lab has previously described the use of an injectable hydrogel depot technology for sustained delivery of vaccine components to improve humoral immune response to infectious diseases such as influenza and SARS-CoV-2.^[21–25]^ Vaccines delivered with these polymer-nanoparticle (PNP) hydrogels elicit lower levels of systemic cytokines often associated with poor tolerability.^[23]^ In this work, we sought to leverage the PNP hydrogel technology to evaluate whether the sustained delivery of a shingles subunit vaccine would overcome the reduced immune responses typically associated with immunosenescence in aged mice (**Figure 1**). We measured humoral and cellular immune responses in young and aged mice and demonstrated that while a PNP hydrogel-based vaccine induced similar responses to an AS01-adjuvanted vaccine in young mice, the sustained-release hydrogel vaccine exhibited more potent and durable responses in aged mice. Since cell-mediated immunity is recognized as the strongest immunological predictor of protection against herpes zoster,^[9, 26–29]^ we quantified T cell responses in young and aged mice and demonstrated negligible age-related changes in responses with PNP hydrogel-based vaccines. Moreover, analysis of serum cytokines associated with systemic reactogenicity confirmed the tolerability of these PNP hydrogel-based vaccines. Overall, we report a simple and effective vaccination strategy leveraging sustained release to improve responses against herpes zoster in aged populations.

**Figure 1.**
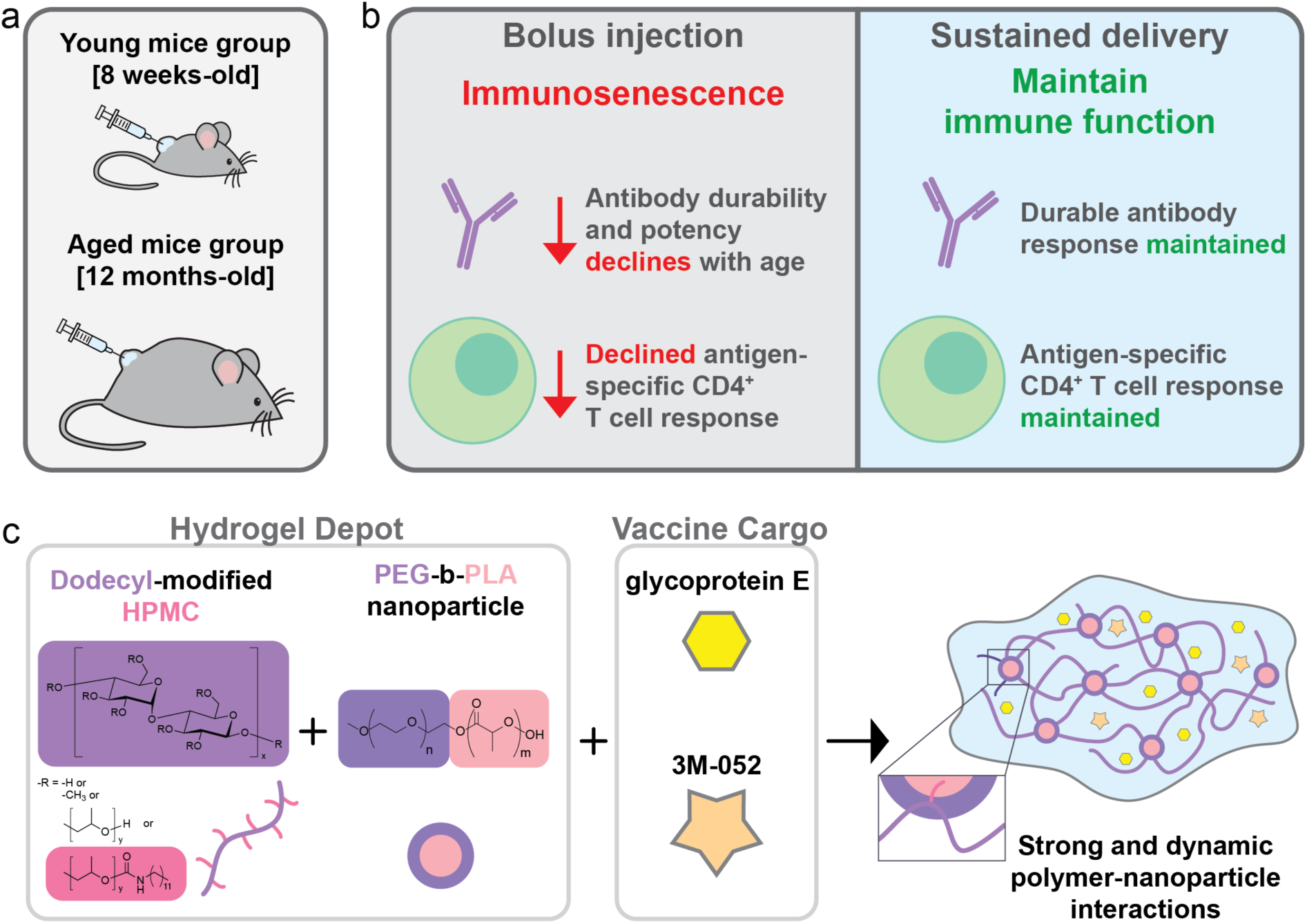
Polymer-nanoparticle (PNP) hydrogel-based shingles vaccine. (a/b) Sustained delivery of vaccine components enables maintenance of durable gE-specific antibody response and CD4^+^ T cell response with age in mice. (c) Schematic of formulation of PNP hydrogels containing vaccine cargo comprising gE antigen and 3M-052, a potent TLR7/8 agonist. Dynamic, multivalent interactions between the hydrogel components yield injectable hydrogel networks that form robust depots for sustained cargo delivery.

## 2. Results

### 2.1 Hydrogels for Slow Delivery of Shingles Vaccine

We have previously demonstrated that prolonged delivery of vaccine antigens and adjuvant molecules with PNP hydrogels can improve humoral immune responses by better mimicking the natural infection timeline.^[22, 24, 30]^ These hydrogels are formulated by mixing dodecyl-modified hydroxypropylmethylcellulose (HPMC-C_12_) with biodegradable nanoparticles (NP) formed with poly(ethylene glycol)-b-poly(lactic acid) (PEG-PLA) (**Figure 1b**). By simply mixing aqueous solutions of HPMC-C_12_ polymers and PEG-PLA NPs, the interaction between the two components forms dynamic, entropy-driven multivalent crosslinking that generates robust physically-crosslinked hydrogels that are injectable and self-healing.^[31]^ Vaccine cargo of interest can be premixed into the NP solution and encapsulated in the hydrogel network.^[32]^ PNP hydrogels display unique temperature independent mechanical properties of shear-thinning and self-healing behavior, enabling the injection of preformed hydrogel through needles and the depot formation in the subcutaneous space.^[21, 33, 34]^ The hydrogel mesh size and retention time *in vivo* can be precisely engineered by varying the concentration of each component, thereby allowing prolonged delivery over controlled timescales.^[24, 30, 34, 35]^ In this work, we chose a PNP hydrogel formulation consisting of 2 wt% of HPMC-C_12_ polymer and 10 wt% of PEG-PLA NPs (denoted PNP-2-10) for further analysis. PNP-2-10 was reported to co-encapsulate and release components of various molecular sizes at a comparable rate with almost complete retention when tested *in vitro*. This hydrogel depot is also known to gradually dissolve *in vivo* and release cargo by erosion, allowing sustained release of entrapped cargo over 4 weeks.^[21–25, 30, 34, 35]^ We selected 3M-052 (Solventum), a lipidated imidazoquinoline derivative and potent TLR 7/8 agonist adjuvant, to encapsulate alongside gE protein in PNP gels based on our previous reports showing enhanced humoral responses with minimal reactogenicity. We hypothesized PNP-2-10 formulations would ensure co-encapsulation and similar release kinetics of gE protein (∼60 kDa) and 3M-052 (lipidated small molecule) based on the hydrogel erosion, given previous reports on almost complete retention of other protein cargo of relevant size and 3M-052 in these formulations.^[21]^

### 2.2 Rheological characterization of PNP 2-10 hydrogel

To confirm that critical rheological properties of the PNP-2-10 hydrogels relating to the injectability and depot formation are maintained with the inclusion of gE protein and 3M-052, we performed various shear rheometry tests (**Figure 2**). PNP hydrogel-based vaccines undergo three administration regimes during use: (i) formulation of the hydrogels with vaccine cargo encapsulated, (ii) injection through a clinically relevant sized needle, and (iii) instant recovery of hydrogel network for retention of cargo post injection and depot formation.^[30, 36]^ We first assessed frequency sweep measurements to demonstrate that vaccine-containing PNP-2-10 formulations exhibit robust gel-like behavior. Vaccine-loaded PNP-2-10 hydrogel was formulated by dissolving gE and 3M-052 in phosphate buffered saline with PEG-PLA NPs and loading into a syringe (**Figure 2a**, shown in blue) and mixing with an HPMC-C_12_ solution loaded into another syringe (**Figure 2a**, shown in opaque white). We observed that over the entire frequency regime tested, vaccine-loaded PNP-2-10 showed higher storage modulus (G′) than loss modulus (G″), confirming solid-like hydrogel formation in the presence of the vaccine components (**Figure 2b**). Steady-state flow rheology elucidates shear-thinning behavior of vaccine-loaded PNP-2-10 with a high shear-thinning index of 0.98, confirming injectability (**Figure 2c and 2d**).^[37]^ Upon injection, once the high shear rates are removed, the hydrogel network quickly reestablishes and the gels regain their solid-like behavior, which is necessary for rapid formation of a depot to minimize burst release and ensure prolonged delivery (**Figure 2e and 2g**).^[34, 38]^ Step-shear measurements in alternating cycles of high shear (10 s^-1^) and low shear (0.1 s^-1^) rates were conducted to ensure the hydrogel’s ability to shear-thin and self-heal (**Figure 2f**). Instant decrease and recovery of viscosity were detected upon applying and removing high shear, confirming rapid network self-healing. We measured the apparent yield stress of vaccine-loaded PNP-2-10 hydrogel to be 159 Pa, which has previously been shown to be sufficient for formation of a robust depot following transcutaneous injection into the subcutaneous space (**Figure 2h**). Similar to our previous reports, the addition of vaccine components did not interfere with the mechanical properties of the PNP hydrogels.^[21–23]^

**Figure 2.**
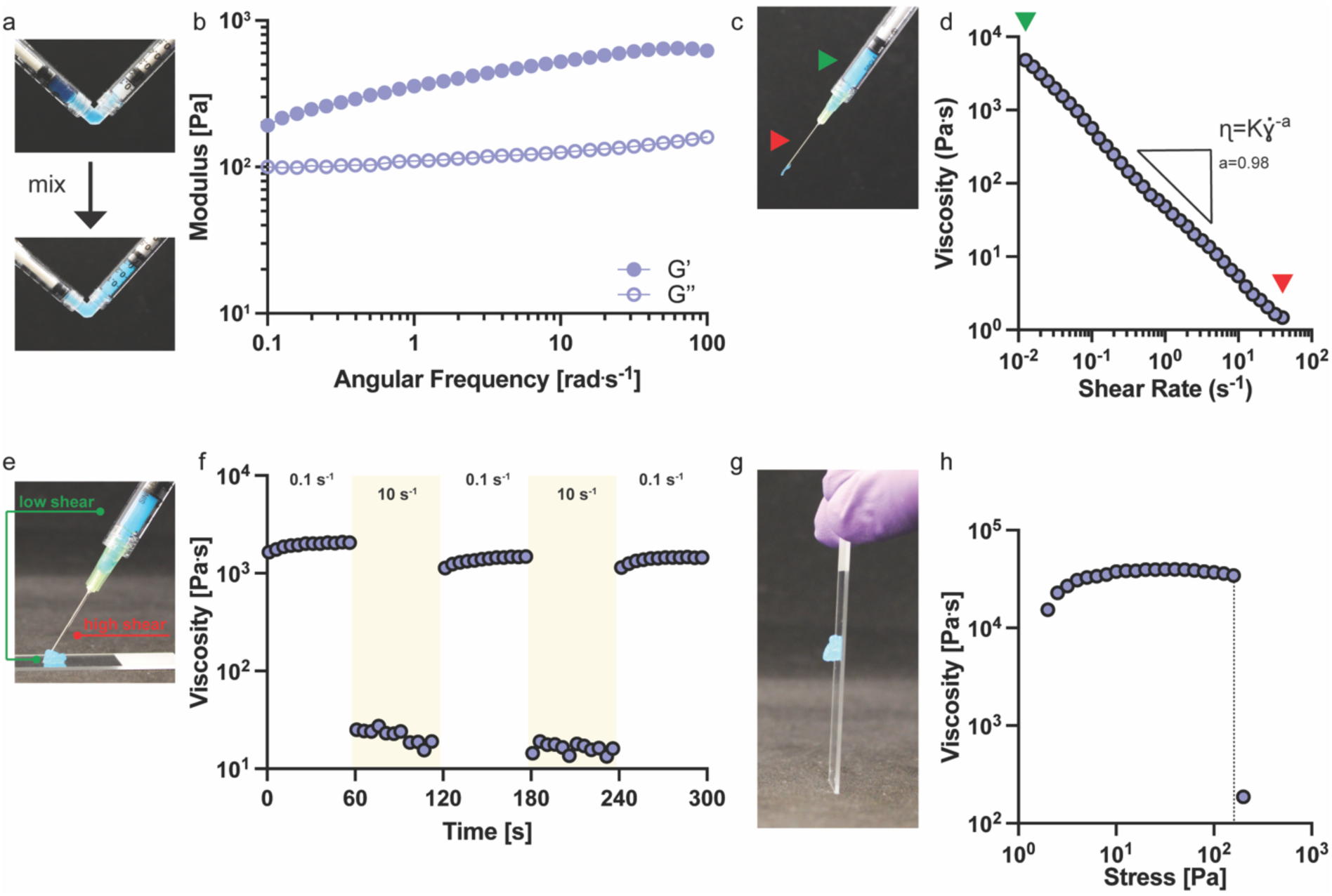
Rheological characterization of vaccine-loaded PNP hydrogel. (a) Representative image of the PNP hydrogel formulation process whereby one syringe is loaded with HPMC-C_12_ polymer solution (opaque white), a second syringe is loaded with PEG-PLA NP solution containing the vaccine components (blue color included for visual comparison), and the two solutions mixed. (b) Frequency sweep of vaccine loaded PNP-2-10 hydrogel at 1% strain. (c) Injection of PNP-2-10 hydrogel through 21-gauge needle. The green arrows indicate the low shear regime while the red arrow indicates the high shear regime. (d) Flow sweep from low to high shear rates for vaccine-loaded PNP-2-10 hydrogels. (e) Instant self-healing of PNP-2-10 hydrogels is displayed post-injection. (f) Step-shear measurements alternating between high (10 s^-1^) and low (0.1 s^-1^) shear rates applied in three cycles to confirm the decrease in viscosity upon application of shear (shaded in yellow) and rapid recovery upon removal of shear. (g) PNP-2-10 hydrogel forms a robust depot post-injection. (h) Stress-controlled flow sweeps measured the apparent yield stress of PNP-2-10 hydrogel.

### 2.3 3M-052 Adjuvanted Hydrogel Vaccine Promotes Robust and Durable Antibody Response with Reduced Inflammation in Aged Mice

We then evaluated the immune response induced by PNP hydrogel vaccines both in young (8-weeks-old; C57BL/6) and aged mice (12-months-old; C57BL/6). These aged mice have been shown to have immune systems roughly comparable to 50 years old humans, which is the recommended age to start receiving shingles vaccine according to CDC.^[39–42]^ We hypothesized that sustained vaccine exposure with 3M-052-adjuvanted hydrogels would improve antigen-specific immune responses and mitigate immunosenescence, while also reducing reactogenicity, compared to the commercial Shingrix® subunit shingles vaccine. Therefore, mice were immunized subcutaneously with either 100μl of a bolus control vaccine comprising gE protein (5μg) and AS01 adjuvant (5μg) as a control to mimic the current clinical vaccine, or 100μl of PNP hydrogel vaccines containing gE protein (5μg) and 3M-052 (1μg) (**Figure 3a**). We also assessed a bolus control vaccine comprising 3M-052/Alum (AAHI), a clinically relevant adjuvant, to independently gauge the impact of the sustained vaccine release from PNP-2-10 hydrogels (**Figure S1-S3**). The dosages of gE and AS01 were selected to match previous reports on human to mouse dosage conversion (1/10^th^ of the human dosage),^[9, 43]^ and 3M-052/Alum dose was chosen based on previous reports.^[21, 22, 44]^ Prime-boost vaccines were delivered on week 0 and week 8, and serum was collected weekly to quantify total IgG antibody titers. The boosting timeframe was chosen to replicate the regimen for the Shingrix® vaccine.^[39]^

**Figure 3.**
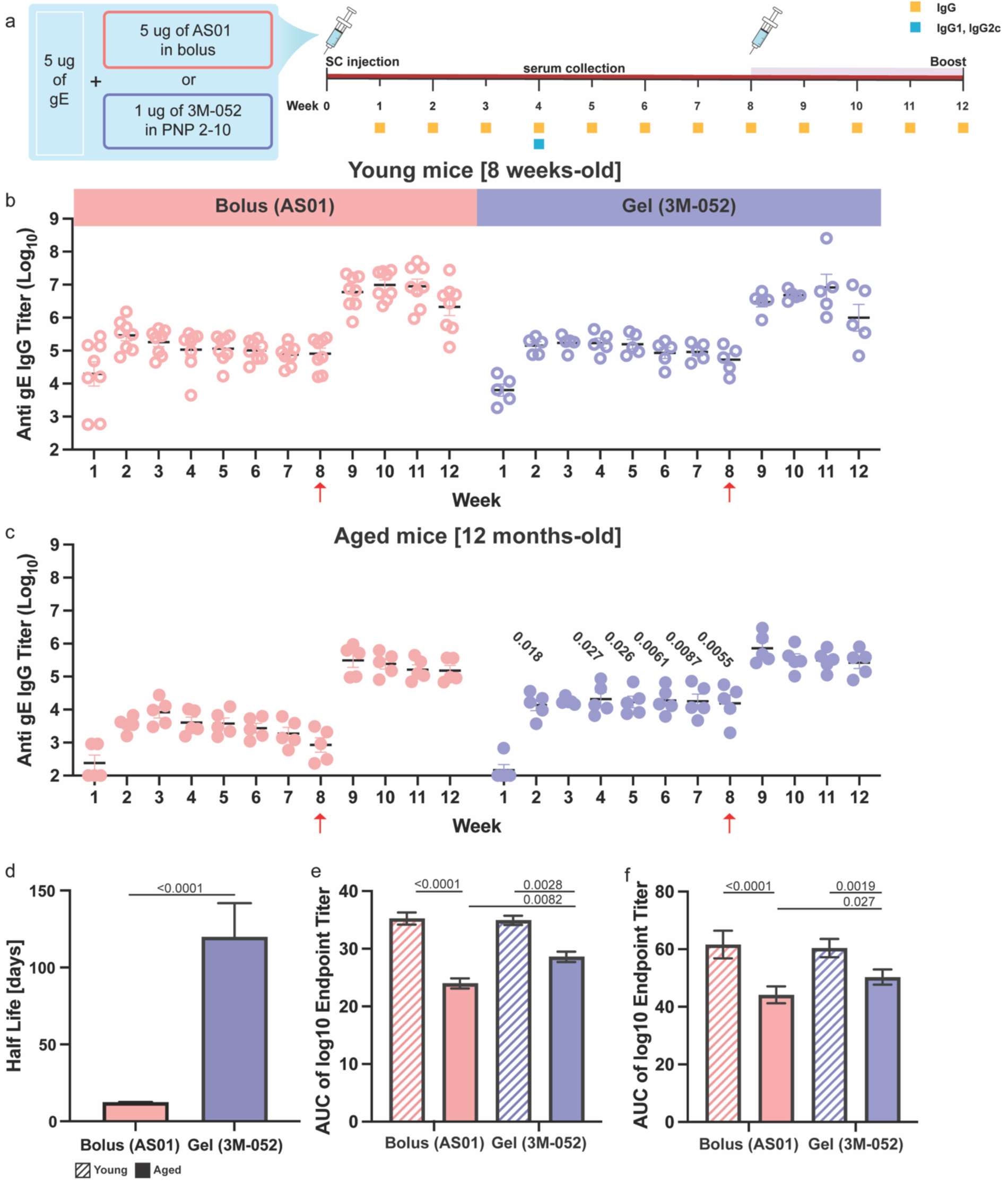
*In vivo* humoral response to gE subunit vaccine. (a) Vaccination and monitoring schedule. Mice were vaccinated with 5μg of gE and 5μg of AS01 for the bolus group or 1μg of 3M-052 for the hydrogel group at week 0 and week 8, and assays were performed as indicated. (b) Anti-gE IgG endpoint titers for young (8-week-old) mice treatment groups (n=8 for the bolus group and n=5 for the hydrogel group). (c) Anti-gE IgG endpoint titers for aged (12-month-old) mice treatment groups (n=5 for both bolus and hydrogel groups). (d) Half-life of antibody decay from week 3 to 8 from (c) in aged mice group. (e) Area under the curve from week 0 to 8 from (b) and (c). (f) Area under the curve from week 0 to 12 from (b) and (c). One-way ANOVA was used for multiple comparisons, and only p-values below 0.05 are displayed.

Young mice treated with 3M-052-adjuvanted PNP-2-10 hydrogel vaccines exhibited total anti-gE IgG endpoint titers comparable to those of the bolus vaccine with AS01, both post-prime and post-boost (**Figure 3b**). Given that AS01 is one of the most potent adjuvants used in modern vaccines and has been shown to induce strong humoral responses when administered with recombinant gE, it is remarkable that the PNP hydrogel vaccine effectively elicited comparable antibody responses.^[9, 13, 45]^ Interestingly, the bolus 3M-052/Alum group produced antibody titers around 200-fold lower than the AS01 and 3M-052-adjuvanted PNP-2-10 hydrogel vaccines (**Figure S1**), as well as two non-responding animals. These observations confirm that sustained vaccine delivery with the PNP hydrogel depot technology is important for inducing stronger antibody responses.

We then analyzed the total IgG antibody titers against gE for the bolus and hydrogel groups in aged mice (**Figure 3c**). Aged mice vaccinated with the AS01 bolus control vaccine elicited slower seroconversion and delayed antibody responses (peak responses observed at week 2 post-injection in young mice and week 3 in aged mice), and 36-fold lower peak antibody responses post-prime compared to young mice. In contrast, the decrease in titer response by age was much less pronounced (8-fold decrease) in the hydrogel group. Notably, while the titers declined rapidly after the post-prime peak in aged mice receiving the bolus AS01 vaccine, the durability of the post-prime titers is remarkably improved in aged mice receiving the hydrogel vaccine. Indeed, there was a 20-fold increase in antibody titers in the hydrogel group compared to the AS01 bolus (p=0.0055). The titers between the bolus group and hydrogel group became comparable after the boost. A similar trend was observed for the bolus 3M-052/Alum group in aged mice whereby the potency and durability of the antibody titers were poor compared to the AS01 or 3M-052 hydrogel groups (**Figure S1 and S2**). In summary, we confirmed that sustained vaccine delivery with the PNP hydrogel induces humoral immune responses comparable to or exceeding that of the clinical standard AS01-based vaccines.

To quantify antibody durability, we applied an exponential decay fit model with bootstrapping to titer data after the peak (week 3) to estimate the half-life of anti-gE antibody titers in aged mice (**Figure 3d**). The half-life of the antibody decay in the hydrogel group was significantly longer at ∼120 days compared to the bolus group at ∼12 days (p<0.0001). Quantification of the area-under-the-curve (AUC) of the antibody response following the prime injection reflected this same pattern (**Figure 3e**). While the hydrogel group elicited comparable antibody titers to the bolus AS01 group in young mice, the hydrogel group elicited markedly enhanced antibody titers compared to the bolus group in aged mice (1.2-fold increase; p=0.0082). A similar trend was observed in AUC including the post-boost titers, with the hydrogel group showing a significant increase over the bolus AS01 group (1.13-fold increase; p=0.027) in aged mice, though the effect was less pronounced given that the titers became comparable post-boost (**Figure 3f**). Again, sustained vaccine exposure with the PNP hydrogel depot was necessary to observe this improvement in humoral immune responses given that the bolus 3M052/Alum group failed to elicit comparable antibody titers to the AS01 adjuvant (**Figure S3**). For this reason, hereafter we limited our comparisons to the clinical standard AS01-adjuvanted control vaccine group. Overall, these results demonstrated that the PNP hydrogel vaccine system comprising the 3M-052 adjuvant mitigates the decay in humoral immunity observed in aged mice compared to clinical standard AS01-adjuvanted control vaccine.

We then quantified subclasses of IgG at week 4 to evaluate the influence of the AS01 and 3M-052-adjuvanted hydrogels on immune signaling. Specifically, we evaluated IgG1 and IgG2c isotypes since these subclasses serve as indirect indicators for Th-2 and Th-1 skewed immune response, respectively.^[46]^ IgG1 titers were not significantly different across all groups in either young or aged mice (**Figure 4a**). In contrast, we observed a trend towards decreasing IgG2c titers with age in both the bolus AS01 and the 3M-052/PNP-2-10 groups (**Figure 4b**). Noticeably, the hydrogel vaccinated groups displayed higher IgG2c titers compared to the bolus AS01 controls. Indeed, the hydrogel vaccine elicited roughly 2-fold higher IgG2c titers in aged mice (3.3×10^4^) than the AS01 vaccine elicited in young mice (1.58×10^4^). The enhanced IgG2c titers in the hydrogel groups resulted in a more balanced Th1/Th2 response than the AS01-based vaccines, with a IgG2c/IgG1 ratio close to 1 for both young and aged mice (**Figure 4c**).

**Figure 4.**
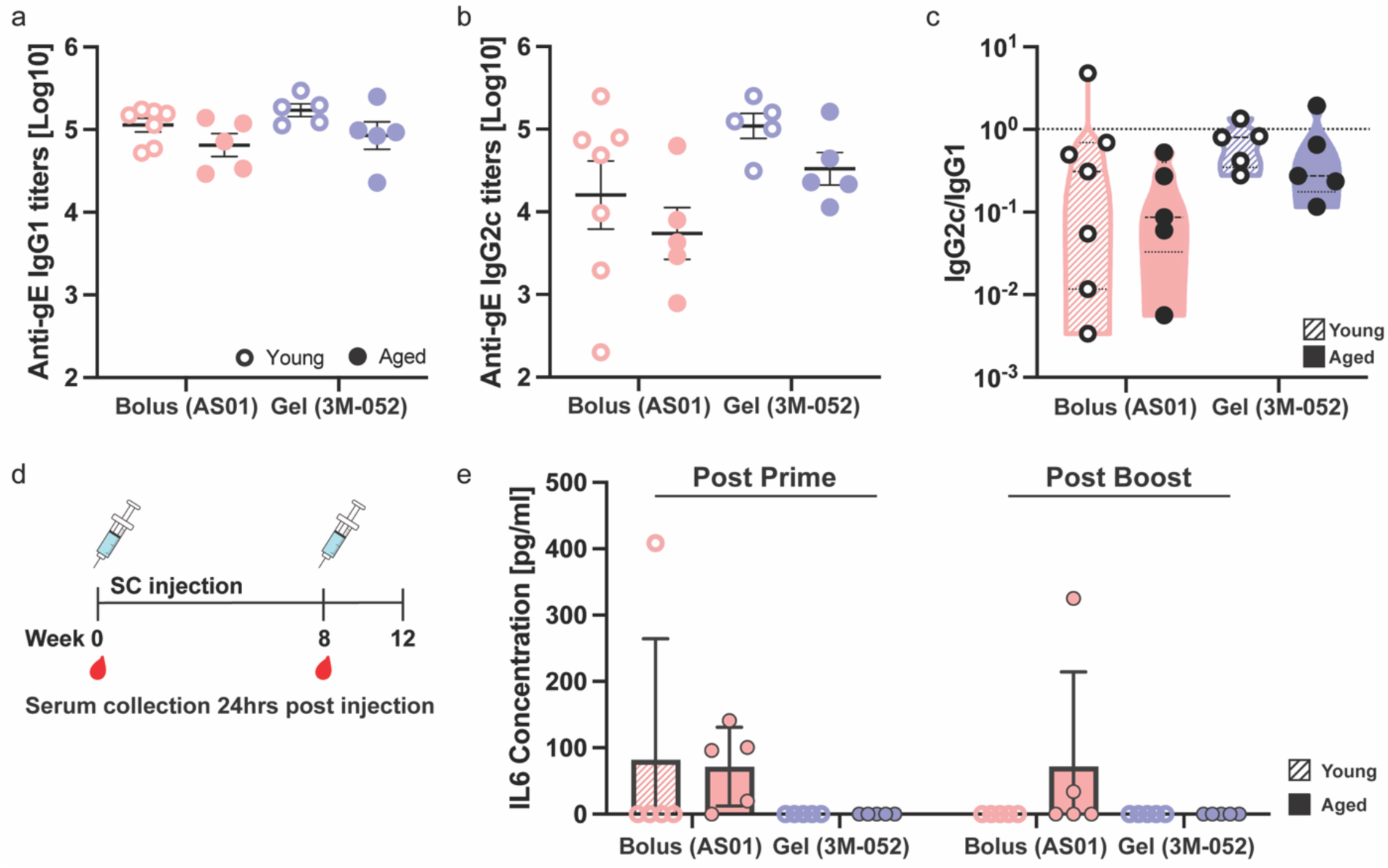
Antibody subtype response and circulating inflammatory cytokine response to gE subunit vaccine. (a) Anti-gE IgG1 and (b) IgG2c titers at week 4. (b) Ratio of anti-gE IgG2c to IgG1 at week 4. Ratio below 1 indicates more Th2 skewed response. (d) Timepoint of serum collection for circulating cytokine measurement. All serum samples were collected 24 hours after the injection. (e) IL6 concentration of young and aged mice 24 hours after injection at week 0 and 8. One-way ANOVA was used for multiple comparisons, and only p-values below 0.05 are displayed.

Interleukin-6 (IL6) is a cytokine that is one of the more strongly correlated markers for reactogenicity.^[12]^ To evaluate the reactogenicity associated with our immunizations, we measured serum IL6 levels after prime and boost administrations (**Figure 4d**). Sera were collected 24 hours after injection. While only one of the young mice treated with the AS01-based vaccine exhibited elevated IL6 concentrations after the prime administration (**Figure 4e**), four of the five aged mice receiving the AS01-based vaccine exhibited elevated IL6 concentrations post-prime. Following the boost administration, elevated IL6 levels were detected only in aged mice receiving the AS01-based vaccine. In contrast, none of the animals receiving the 3M-052-adjuvanted hydrogel vaccines exhibited elevated IL6 levels, suggesting that hydrogel vaccines were well tolerated in both young and aged cohorts, consistent with previous reports.^[23, 25]^ These studies indicate that PNP hydrogel vaccines adjuvanted with 3M-052 can mitigate age-related decreases in vaccine potency without increased reactogenicity typically observed with the commercial AS01-based vaccine.

### 2.4 Antigen specific CD4^+^ T cell response was improved when immunized with 3M-052 Adjuvanted Hydrogel Vaccine in aged mice

In addition to humoral immune reponses, maintaining VZV-specific T cell populations is central to preventing symptomatic recurrence and promoting recovery from varicella.^[29, 47]^ Yet, these T cell populations are highly affected by immunosenescence, leading to a decrease in functional VZV-specific T cells with age.^[29, 48, 49]^ Previous studies have shown that 8-week-old mice immunized with recombinant gE protein and AS01 adjuvant displayed enhanced gE-specific functional CD4^+^ T cell responses and the prevalence of these cells is a strong indicator of vaccine efficacy in controlling VZV replication during reactivation.^[9, 50, 51]^ We therefore analyzed antign-specific interfeuron-γ (IFN-γ) producing CD4^+^ T cells to characterize the cellular response induced in young and aged mice groups by our hydrogel-based vaccines and bolus controls. Spleens were explanted 4 weeks after the boost immunization and stimulated with VZV gE peptides for ELISpot analysis (**Figure 5a**). Young mice immunized with the 3M052-adjuvanted hydrogel vaccine induced comparable IFN-γ^+^ CD4^+^ T cell responses compared to that of the AS01-based bolus control vaccine (**Figure 5b**). While a significant decline in these cellular responses with age was evident in the bolus group (13-fold decrease; p=0.032), a negligible difference by age was observed in the hydrogel group (1.7-fold decrease; p=0.99) (**Figure 5c and 5d**). These observations substantiate that sustained vaccine exposure with the PNP hydrogel depots adjuvanted with 3M-052 attenuate age-related decline in cell-mediated immunity by enhancing CD4^+^ T cell responses in aged animals.

**Figure 5.**
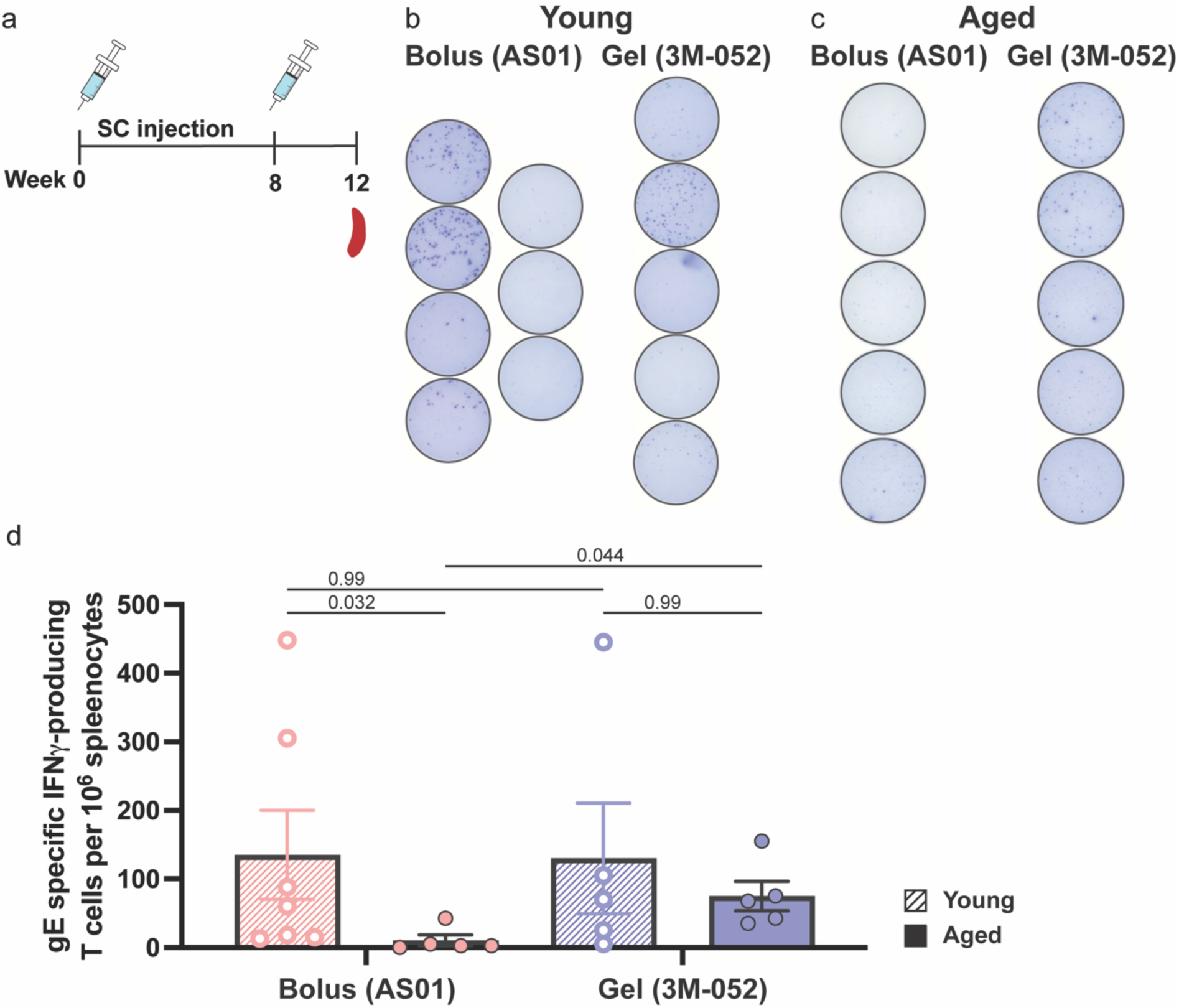
gE-specific CD4^+^ T cell responses measured by ELISpot. (a) Experiment timeline of injection schedule and spleen collection at week 12. (b) Images of visible spots from IFNγ producing CD4^+^ T cell populations in young mice groups and (c) aged mice groups (d) IFNγ producing CD4^+^ T cell counts per 10^6^ splenocytes per well upon stimulation with gE. One-way ANOVA was used for multiple comparisons. Data were log-transformed prior to statistical analyses to normalize variance and meet requirements for statistical tests used in (d).

### 2.5 Robust germinal center responses were observed in the PNP hydrogel vaccinated group in the draining lymph node

To better understand the mechanism of action of these sustained-release vaccines, we evaluated the germinal center activity in the draining lymph node two weeks after prime injection in aged mice using flow cytometry (**Figure 6a** and **Figure S4**). While the total number of lymphocytes in both vaccine groups were comparable, we detected significantly higher frequency and percentages of germinal center B cells (GCBCs) in the groups vaccinated with hydrogel-based vaccines (**Figure 6b-e**). We further quantified gE protein specific GCBCs and observed a similar trend, whereby significantly higher counts and percentages of antigen specific-GCBCs were observed in the hydrogel group compared to the bolus AS01-based control (**Figure 6f-h**). We then measured T follicular helper (T_FH_) cells as they are important in the formation of germinal centers and maintenance of GCBCs.^[52, 53]^ We observed a trend towards higher counts and percentages of T_FH_ cells in the groups receiving the 3M-052-adjuvanted PNP-2-10 hydrogel vaccines compared to the AS01 bolus control, although these differences were not statistically significant (**Figure 6j-k**). Similarly, the number of plasma cells in the GCs was comparable in bolus and hydrogel groups (**Figure 6l–m**). Additionally, we evaluated the ratio of GCBCs to T_FH_ as a metric for the quality of T_FH_ help,^[54, 55]^ which demonstrated that groups immunized with the hydrogel-based vaccines exhibited a trend towards a higher ratio of GCBCs:T_FH_ than the bolus AS01-based vaccines (**Figure 6n**). With aging, a reduction in somatic hypermutation as well as a decreased population of plasma cells, GCBCs, and antigen-specific GCBCs is observed, possibly due to decreased functionality of T_FH_ associated with age.^[56–61]^ Importantly, sustained vaccine exposure in the hydrogel-based vaccine groups showed promise in improving both GCBC and antigen-specific GCBC responses, and suggested a tendency towards enhanced T_FH_ prevalence and function compared to the clinical control. Overall, these experiments suggest that sustained vaccine exposure with the PNP-2-10 hydrogel depot technology can mitigate age-related decreases in GC activity.

**Figure 6.**
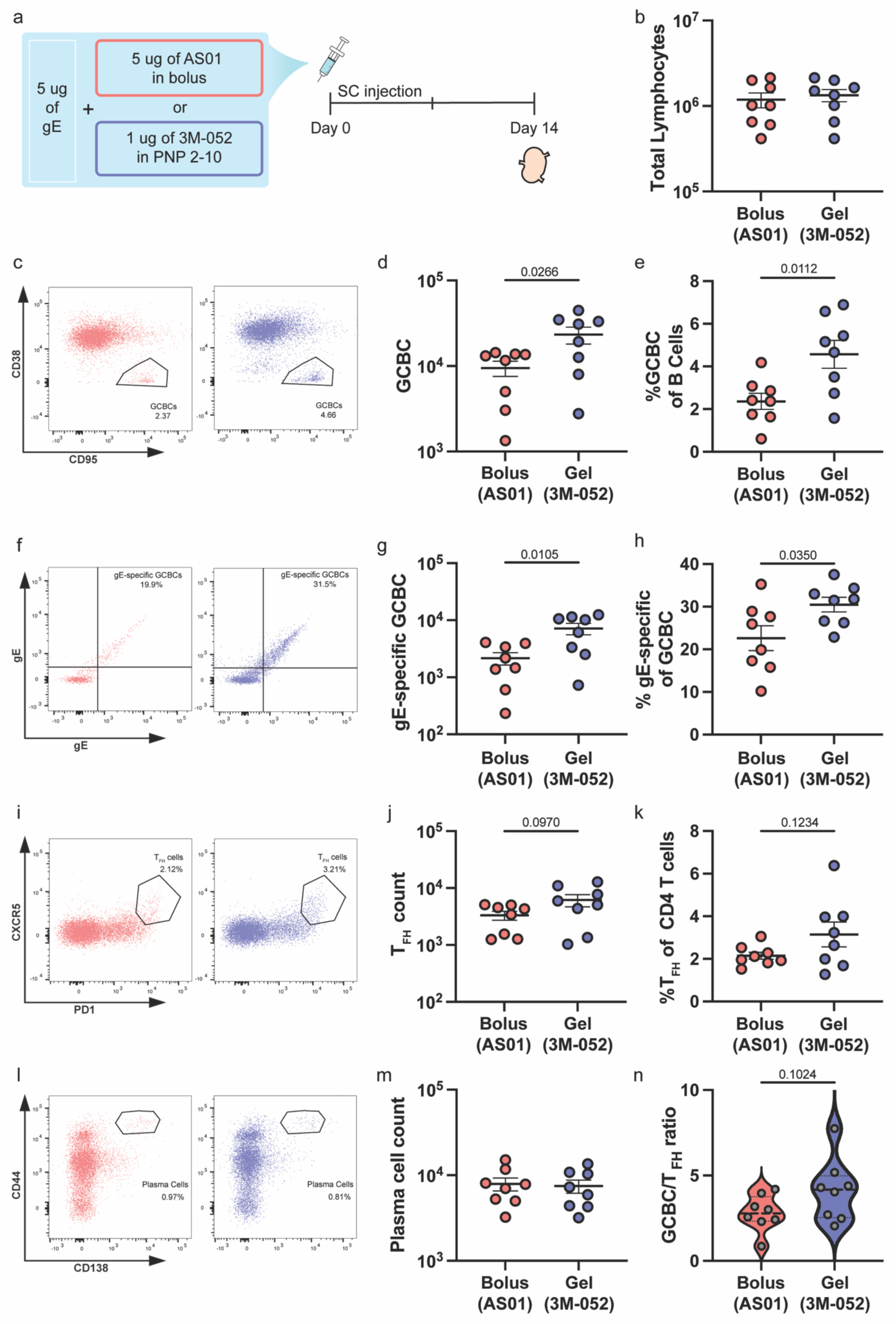
Cytometry analyses of germinal center responses. (a) Immunization and lymph node collection schedule. Mice were immunized with 5μg of gE in bolus with 5μg of AS01 or in hydrogel with 1μg of 3M-052 at week 0. Lymph nodes were harvested at week 2. (b) Count of total lymphocytes. (c) Representative flow plot of B cell population and quantification of (d) counts and (e) percentage of GCBC of all B cells. (f) Representative flow plot of GCBC population and quantification of (g) counts and (h) percentage of antigen-specific GCBC. (i) Representative flow plot of CD4^+^ T cells population and quantification of (j) counts and (k) percentage of T_FH_ of all CD4^+^ T cells. (l) Representative flow plot of live cell population and quantification of (m) counts of plasma cells of all live cells. (n) Ratio of GCBC to T_FH_ cells. One-way ANOVA was used for multiple comparisons.

## 3. Discussion

The global aging population is more susceptible to emerging infectious diseases, as shown in the recent outbreak of COVID-19, and exhibits increased morbidity and mortality from established pathogens such as influenza, which remains particularly lethal among individuals aged over 60 years.^[62–65]^ Therefore, the ability to enhance vaccine efficacy in this demographic remains a critical and pressing healthcare challenge. Current approaches to improving immune responses in the elderly include optimizing vaccine design through modification of the administration route, dosing regimen, vaccine dosing, and incorporation of novel adjuvants.^[8, 66]^ While these strategies have shown promising results for some vaccines, exemplified by the success of inclusion of the AS01 adjuvant in the Shingrix® vaccine, any accompanying reactogenicity typically leads to reduced vaccine compliance.^[16]^ Herein, we report the use of an injectable hydrogel depot technology as a delivery platform to prolong the co-delivery of shingles vaccine components to improved humoral and cellular immune responses in aged populations without reactogenicity. We show that both the humoral and cell-mediated immunity are improved in the aged mouse model compared to the current clinical vaccine, both of which are important determinants of determining the success of the shingles vaccine. Our hydrogel vaccine is well tolerated with no detectable inflammatory cytokine associated with reactogenicity. Importantly, we emphasize that because the PNP hydrogel can enhance vaccine response, it enables the use of 3M-052, a less reactogenic but less immunogenic adjuvant 3M- 052. We show that 3M-052 administered as a bolus vaccine adjuvant with alum resulted in poor or no response, whereas in the PNP hydrogel, we observe a comparable or better humoral and cell mediated responses compared to AS01 bolus without reactogenicity.

We have previously reported that the use of PNP hydrogel enhances the breath, potency, and durability of the antibody response following a single administration. We also demonstrated the tolerability of using PNP hydrogel as a sustained vaccine delivery platform. We hypothesized that a hydrogel-based vaccine delivery method could mitigate the effect of immunosenescence, which leads to reduced antibody response. Indeed, we identified a durable and potent antibody response induced in aged mice vaccinated with PNP hydrogel. Furthermore, we demonstrated that IL6 levels in hydrogel vaccinated groups were below the detection limit, whereas groups vaccinated with AS01-adjuvanted bolus exhibited an age-associated increase in IL6 levels. These results demonstrate that sustained delivery technique with PNP hydrogel improves the durability and potency of humoral response while reducing the reactogenicity associated with the current clinical vaccine.

Cell-mediated immunity to VZV is identified as a key immunological predictor of reduced herpes zoster incidence, severity, and postherpetic neuralgia according to clinical investigations.^[26–28]^ Studies suggest that VZV-seropositive young adults exhibit high levels of circulating gE-specific memory CD4+ T cells, which help to control the viral replication.^[51]^ These individuals also retain relatively high level of VZV-specific T cell responses over time, which corresponds to the low herpes zoster incidence.^[47, 67]^ With aging, as both the innate and adaptive immune system decline in function, decline in VZV-specific T cell population is observed.^[26, 68]^ Hence, a vaccine that enhances VZV-specific cell-mediated immunity in individuals with decreased immunity should offer protection against herpes zoster. Our previous works on PNP hydrogel vaccine mainly focused on the enhanced humoral responses while cell-mediated immunity was less explored.^[21–25, 30]^ We observed antigen-specific CD4+ T cell response comparable to AS01 adjuvanted soluble formulation. Importantly, the reduction in CD4+ T cells seen in the soluble group with age was not observed in the hydrogel-vaccinated groups with age. These findings indicate that the PNP hydrogel vaccine may counteract age-related immune decline and can improve the vaccine efficacy.

Improving vaccine strategies for the elderly requires enhancing immune response while minimizing inflammation to reduce vaccine hesitancy.^[16, 49, 69]^ We demonstrate that the PNP hydrogel improves both humoral and cellular immune response without age-related decline or increased inflammation. A single administration of the hydrogel vaccine produced a potent and durable antibody response, suggesting that use of our technology may also be effective in improving the efficacy of other vaccines, such as influenza and COVID-19, that have shown reduced protection in the older population. Given that Shingrix is often cited as a benchmark for elderly vaccination, our reporting of comparable or superior results is particularly encouraging. To our knowledge, this work is the first to examine the age-related vaccine efficacy using a sustained delivery technology. These findings highlight the potential of this technology to enhance vaccine effectiveness against infectious diseases that are both more severe and less responsive to vaccination in the elderly.

## 4. Conclusion

In this work, we show that hydrogel-based sustained delivery of shingles subunit vaccine enhances humoral and cellular immune response while minimizing inflammatory responses in aged mice. By counteracting the effects of immunosenescence, this approach mitigates the age-associated decline in antibody response and cell-mediated immunity, key correlates of protection against herpes zoster. The PNP hydrogel formulation adjuvanted with 3M-052 achieved enhanced immunogenicity compared to the clinical standard of AS01-adjuvanted vaccine without associated reactogenicity, suggesting a new pathway toward safer vaccination strategies for the elderly. Given the persistent challenge of suboptimal vaccine efficacy in aging populations, these results underscore the potential of the sustained delivery technique to improve immune protection not only for shingles but also for other infectious diseases, such as influenza and COVID-19 that show age-related reduced vaccine response.

## 5. Materials and methods

### 5.1 Materials

HPMC (US Pharmacopeia Grade), N,N-Diisopropylethylamine (Hunig’s base), hexanes, diethylether, 1-dodecyl isocyanate (99%), N-methyl-2-pyrrolidone (NMP), Poly(ethylene glycol)–methyl ether (PEG, 5 kDa), dichloromethane (DCM), 3,6-dimethyl-1,4-dioxane-2,5-dione (lactide), 1,8-diazabicyclo(5.4.0)undec-7-ene (DBU, 98%), acetonitrile (ACN), dimethyl sulfoxide (DMSO), and bovine serum albumin (BSA) were purchased from Sigma–Aldrich and used as received. 3M-052 was purchased from 3 M and the Access to Advanced Health Institute (AAHI). Zoster vaccine recombinant, adjuvanted Shingrix was purchased from GlaxoSmithKline (GSK) through Stanford Hospital Pharmacy. Goat anti-mouse IgG Fc secondary antibody (A16084) horseradish peroxidase (HRP) was purchased from Invitrogen. 3,3″, 5,5″ - Tetramethylbenzidine (TMB) ELISA substrate, high sensitivity, was purchased from Abcam. Pepmix VZV gE was purchased from JPT peptides.

### 5.2 Preparation of HPMC-C_12_

HPMC-C_12_ was prepared according to previously reported procedures. Briefly, HPMC (1.0g) was dissolved in 40ml of NMP overnight at room temperature. The solution was heated to 50°C for 30 minutes. 125μl of 1-dodecyl isocyanate was dissolved in 5ml of NMP and added dropwise to reaction mixture post 30 minutes of heating with 125μl (∼10 drops) of Hunig’s base (catalyst). The solution was left under heating for 15 more minutes and left overnight at room temperature. The solution was precipitated from acetone and redissolved in water for dialysis for at least 4 days. The polymer was lyophilized and reconstituted with sterile PBS at 6wt% stock solution.

### 5.3 Preparation of PEG-b-PLA

PEG-b-PLA polymer was prepared according to previously reported procedures. Briefly, 2.5g of 5kDa PEG was melted at 100°C and dried on vacuum for at least 2 hrs. 5ml of DCM was used to dissolve PEG and left under nitrogen. 10g of recrystallized, dry lactide was measured in oven-dried 250ml of round flask bottom. Dry lactide was left under vacuum for 5 minutes and flushed with nitrogen for 30 seconds, repeated 3 times to ensure all atmosphere is replaced with nitrogen. 50ml of DCM was added to dissolve lactide. PEG solution was added rapidly to the lactide solution with DBU (500μl of 150ul/ml of DBU in DCM was added). The solution was stirred for 8 minutes before quenching with 500μl of quench solution (3-4 drops of acetic acid in acetone). PEG-b-PLA polymer was collected and dried under vaccum. For quality control (QC) measurement, gel permeation chromatography (GPC) was used to verify the molecular weight and dispersity of polymer.

### 5.4 Preparation of PEG-PLA NPs

PEG-PLA NPs were prepared by adding 1ml solution of PEG-PLA in 75:25 acetomitrile:DMSO mixture (50 mg×ml^-1^) dropwise to 10ml of water under 600rpm of stir rate at room temperature. NPs were centrifuged over a filter (10kDa; Millipore Amicon Ultra-15) at 4500rpm for 1 hr and resuspended in PBS to final concentration of 200 mg×ml^-1^. Dynamic light scattering (DLS) was used to measure NP diameter (32 ± 3 nm).

### 5.5 Preparation of PNP hydrogel

PNP 2-10 hydrogel (2 wt% of HPMC-C_12_ and 10wt% of PEG-PLA NP) was formulated by mixing a 2:3:1 weight ratio of 6 wt% of HPMC-C_12_ polymer solution, 20 wt% of PEG-PLA NP solution, and PBS containing vaccine components of gE subunit protein and 3M-052 adjuvant. HPMC-C_12_ was loaded in one syringe and all the other components, premixed in a 1.5 ml of Eppendorf tube, were loaded in the second syringe. Gels were made by mixing the components in two syringes connected by an elbow connector back and forth for at least 50 times. The elbow connecter was replaced by 21-gauge needle for injection after mixing.

### 5.6 Shear Rheology

Rheological characterization was performed on a TA Instruments Discovery HR-2 stress-controlled rheometer using a 20mm diameter serrated plate with 500μm gap at 25°C. Frequency sweep measurements were performed over frequencies from 0.1 rad^.^s^-1^ to 100 rad^.^s^-1^ (constant 1% strain). Flow sweeps were performed from 100 s^-1^ to 0.1 s^-1^ shear rates. Shear stress sweeps were done from low to high with steady-state sensing to measure yield stress. Yield stress values were defined as the stress at which the viscosity decreases 10% from the maximum. Steady shear experiments were performed by alternating between a 0.1 s^-1^ and 10 s^-1^ for 60 seconds, and the viscosity values were measured every second.

### 5.7 Vaccine Formulations

The antigen dose of 5µg glycoprotein E (gE, GSK) was included in 100µl of bolus or hydrogel vaccine formulation. For bolus clinical control, 5µg of AS01 was included as adjuvant and formulated following manufacturer’s guide. For the alum control group, 100µg of alum and 1µg of 3M-052 were mixed and administered. For PNP hydrogel vaccines, vaccine components were mixed with HPMC-C_12_ and PEG-PLA NP at desired concentrations with 1µg of 3M-052. Mice were boosted on week 8 for all groups with the same vaccine formulation given for the prime injections.

### 5.8 Mice and Vaccination

All animal studies were performed in accordance with the National Institutes of Health guidelines with the approval of Stanford Administrative Panel on Laboratory Animal Care (Protocol APLAC-32109). Eight-week-old for young groups and 12-months-old for aged group of female C57BL/6 mice were purchased from Charles River and housed in the animal facility in Stanford University. Mice were shaved and injected subcutaneously on the right flank with 100µl of bolus or gel vaccine using either 26- or 21-gauge needle, respectively. Blood was collected from the tail vain weekly. Lymph nodes were collected at week 2 for flow cytometry analysis, and spleens were collected for ELISpot analysis at week 12, both after euthanasia under CO_2_. The animal subjects comply with ARRIVE guidelines.

### 5.9 Mouse Serum ELISA

Serum gE-specific IgG antibody titers for the vaccines were measure using an ELISA. 96 well maxisorp plates (Thermofisher) were coated with gE protein (GSK) at 1μg/mL in 1X PBS overnight at 4°C. Between each step, plates were washed 5 times with PBS 1X containing 0.05% of Tween-20. Plates were then blocked with PBS containing 1% non-fat dry milk for 1 h at room temperature. Serum was diluted into a 1% BSA in PBS solution in a v-bottom plate at 1:200 and then four-fold or eight-fold serial dilutions were performed. Titrations were added to plates for 2h at 25°C. Goat– anti-mouse IgG Fc-HRP (1:10,000, Invitrogen, A16084), IgG1 heavy chain HRP (1:20000, abcam ab97240), or IgG2c heavy chain HRP (1:20000, abcam ab97255) was added for 1h at 25°C. Plates were developed with TMB substrate (TMB ELISA Substrate (High Sensitivity), Abcam) for 5 minutes. The reaction was stopped with 1 M HCl. The plates were analyzed using a Synergy H1 Microplate Reader (BioTek Instruments) at 450nm. The total IgG and the subtypes were imported into GraphPad Prism 8.4.1 to determine the endpoint titers by fitting the curves with a five-parameter asymmetrical nonlinear regression at a cutoff of 0.1 (IgG) or 0.2 (IgG subclasses). For cytokine ELISA, mouse IL6 serum cytokine ELISA kit was purchased from R&D systems and used following the manufacturer’s guide.

### 5.10 Half-life Decay Model

Parametric bootstrapping method was used in MATLAB. Titer distribution for each time point was assumed to be log normally distributed, and the fit was applied from week 3 to 8. For each of the 1000 simulations in bootstrap, linear regression was used to fit the sample to an exponential decay model. P-value was determined using a Student’s t-test on GraphPad Prism.

### 5.11 ELISpot

The frequency of gE-specific IFN-γ producing CD4^+^ T cells was measured using the ImmunoSpot Mouse IFN-γ ELISpot Kit from Cellular Technology Limited (CTL). Spleen was harvested on week 12, processed to single cell suspension, and plated at 400,000 splenocytes per well. Spleen cells were restimulated with 1 μg of VZV gE peptide (JPT)/ well for 24 hrs at 37°C. Spots were developed and counted following the manufacturer’s guide.

### 5.12 Flow Cytometry Analysis

Mice were euthanized using carbon dioxide to collect inguinal lymph nodes. After lymph node dissociation into single cell suspensions, staining for viability was performed using Ghost Dye Violet 510 (Tonbo Biosciences, cat. Number 13-0870-T100) for 5 mins on ice and washed with FACS buffer (PBS 1X with 3% FBS, 1mM EDTA). Fc receptors were blocked using anti-CD16/CD32 antibody (clone: 2.4G2, BD Biosciences, cat. Number 553142) for 5 mins on ice and stained with fluorochrome conjugated antibodies: CD19 (PerCP-Cy5.5, clone: 1D3, BioLegend, cat. Number 152406), GL7 (AF488, clone: GL7, BioLegend, cat. Number 144613), CXCR5 (BV711, clone: L138D7, BioLegend, cat. Number 145529), CD4 (BV650, clone: GK1.5, BioLegend, cat. Number 100469), CD138 (BV605, clone: 281-2, BioLegend, cat. Number 142531), CD44 (APC-Cy7, clone: IM7, cat. Number 103028), CD3 (AF700, clone: 17A2, BioLegend, cat. Number 100216), CD95 (PE-Cy7, clone: Jo2, BD Biosciences, cat. Number 557653), PD1 (PE-DazzleTM594, clone: 29F.1A12, BioLegend, cat. Number 135228), anti-gE tetramer (AF647) and anti-gE tetramer (Cy3) for 30 mins on ice. Tetramers were prepared on ice by adding AF647- or Cy3-Streptavidin (Thermo Scientific, cat. Number S21374; ThermoFisher Scientific, cat. Number 434315) to 7.1 μM biotinylated glycoprotein (Sino Biological, cat. 40907-V08H-B) in five steps, once every 20 mins, for a final molar ratio of 5:1 gE protein. After a washing step, cells were analyzed on the Symphony II flow cytometer (BD Biosciences). Data were analyzed with FlowJ 10 (FlowJo, LLC).

### 5.13 Statistics

Comparisons between two groups were conducted using a two-tailed Student’s t-test, whereas comparisons involving more than two groups were done using a one-way ANOVA with Tukey’s post hoc correction for multiple comparisons using GraphPad PRISM. Results with p < 0.05 were considered statistically significant. Data were log-transformed using the formula *y* = log_10_ *x* prior to statistical analyses when necessary to normalize variance and meet requirements for statistical tests used.

## Acknowledgements

This work was supported in part by the Bill & Melinda Gates Foundation (INV027411). Y.E.S. and O.M.S. are thankful for Hancock Fellowship of the Stanford Graduate Fellowship in Science and Engineering. O.M.S., J.Y., and N.E. are thankful for a National Science Foundation Graduate Research Fellowship. B.S.O. is grateful for an Eastman Kodak Fellowship. Cell sorting/flow cytometry analysis for this project was done on instruments in the Stanford Shared FACS Facility (RRID: SCR_017788) using NIH S10 Shared Instrument Grant (1S10OD026831-01). We would also like to thank the staff of the BioE/ChemE Animal Facility who cared for the mice.

## Author Contributions

Y.E.S. and E.A.A. conceived of the idea and designed specific experiments. Y.E.S., J.Y., B.S.O., and O.M.S. performed experiments. Y.E.S and N.E. analyzed the data. Y.E.S. and E.A.A. wrote the paper. J.Y. and B.S.O edited the paper.

## Materials and Data Availability

All data needed to evaluate the conclusions in the paper are present in the paper and/or the Supplementary Materials. Further information and requests for resources or raw data should be directed to and will be fulfilled by the lead contact, Eric Appel.

Received: ((will be filled in by the editorial staff))

Revised: ((will be filled in by the editorial staff))

Published online: ((will be filled in by the editorial staff))

