## Supplemental Data for "Sustained Delivery of a Shingles Subunit Vaccine Overcomes Age-Related Declines in Humoral and Cellular Immunity Relative to Shingrix"

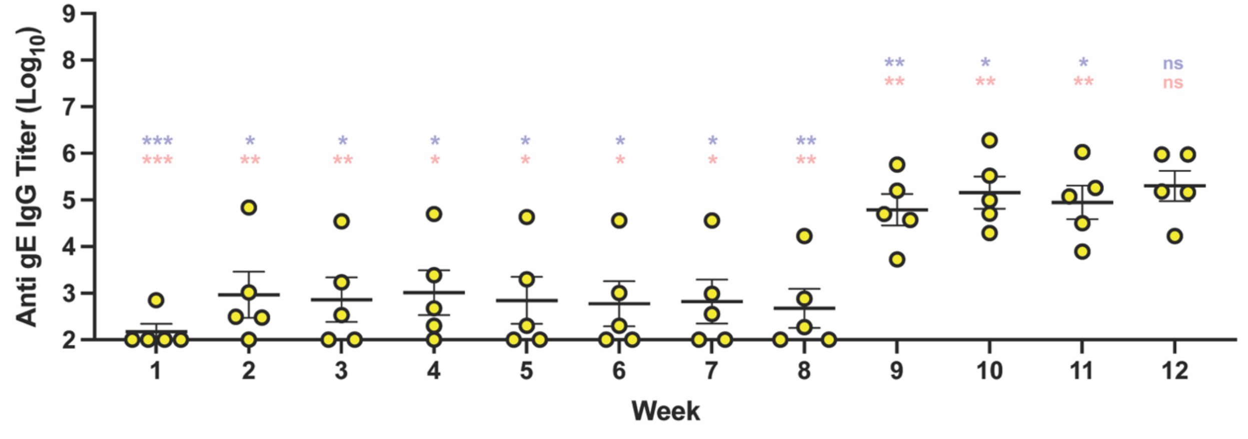


**Fig S1.** Total IgG response to subunit vaccine with alum and 3M-052 in 8-weeks-old mice.

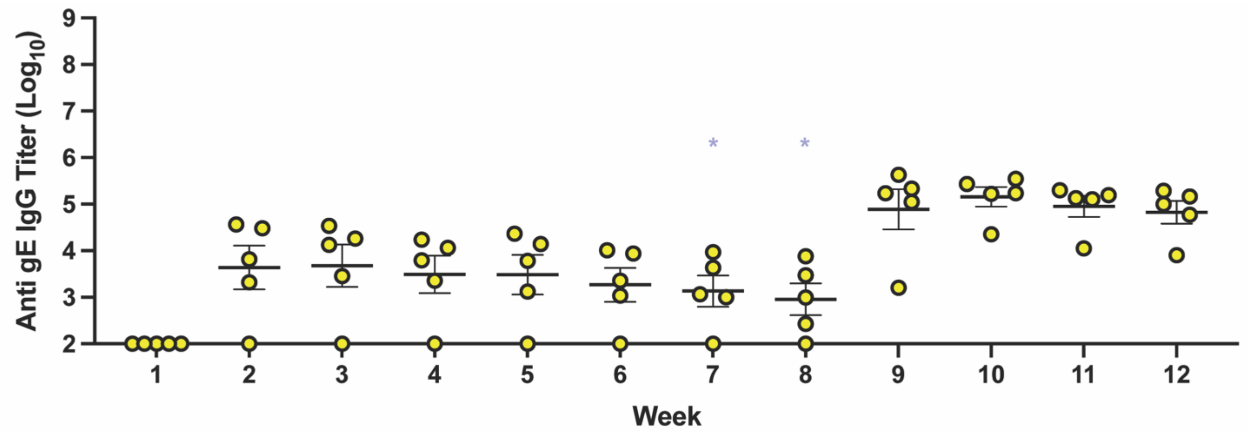


**Fig S2.** Total IgG response to subunit vaccine with alum and 3M-052 in 12-months-old mice.

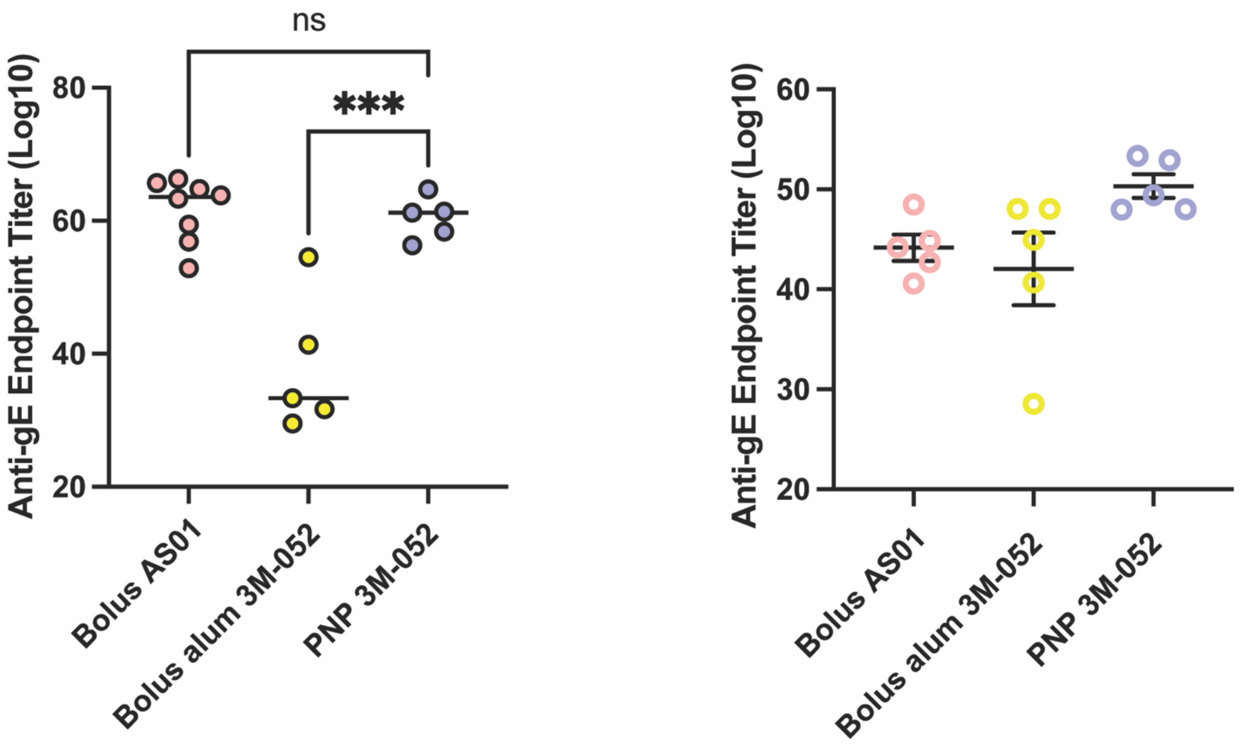


**Fig S3.** AUC from Fig S1 (left) and Fig S2 (right).

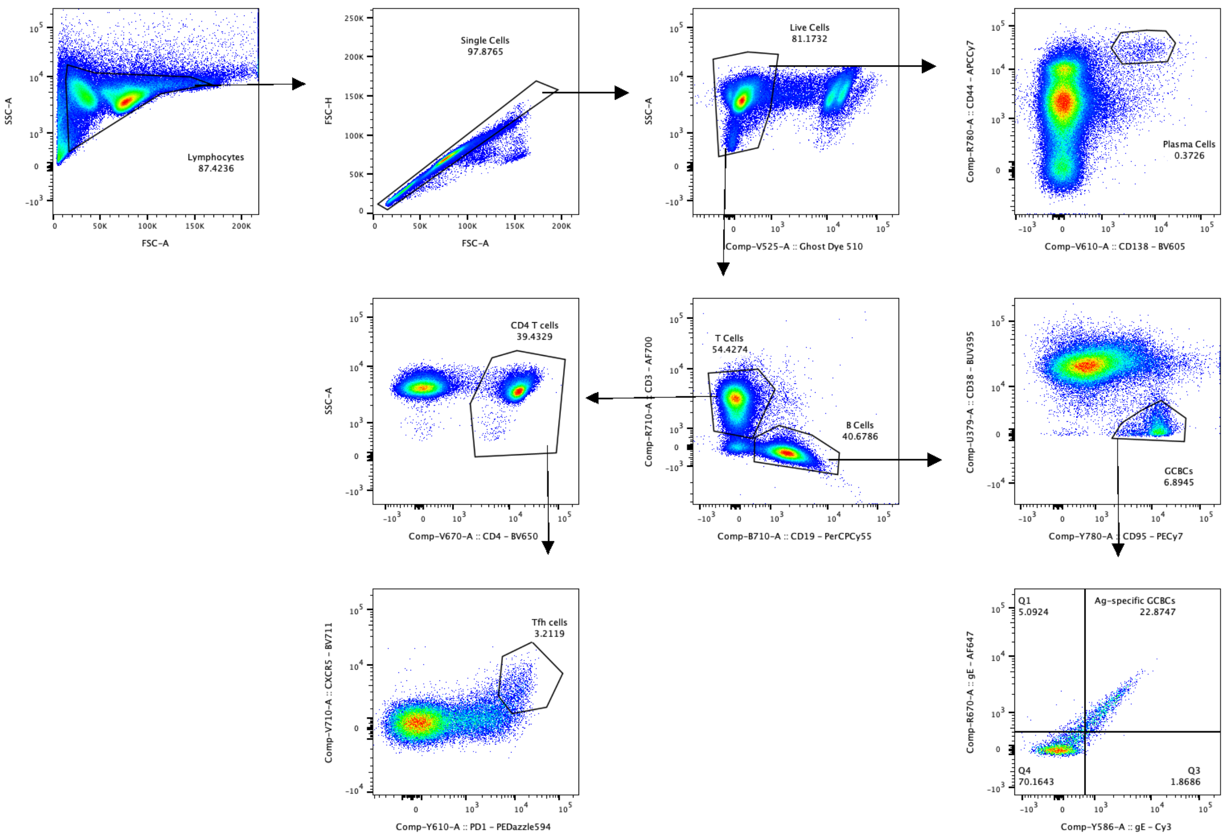


**Fig S4.** Sample gating scheme.
